# Mid-Infrared Photothermal Mesoscopy with Millimeter Field of View and Sub-micron Spatial Resolution

**DOI:** 10.1101/2025.11.03.686410

**Authors:** Rong Tang, Jiaze Yin, Bethany Weinberg, Haonan Lin, Jiahui Lu, David Gate, Oxana Klementieva, Ji-Xin Cheng

## Abstract

By optically sensing mid-infrared absorption through a visible probe beam, mid-infrared photothermal (MIP) microscopy has emerged as a powerful tool for chemical imaging with micromolar sensitivity and submicron spatial resolution. The adoption of spatially multiplexed camera-based widefield detection further enhanced the imaging speed. However, current widefield MIP systems suffer from a small field of view (FOV)—typically tens of square micrometers, which constrains their utility in large-area tissue imaging applications. Here, we report a laser-scan MIP mesoscope that achieves millimeter-scale FOV while preserving submicron resolution. By leveraging an all-reflective laser scanning architecture, low-magnification and medium numerical-aperture objectives, and a defocused signal collection scheme, our MIP mesoscope achieves a 1.2 × 1.2 mm^2^ FOV, 650 nm lateral resolution, and microsecond-scale pixel dwell time. In vivo chemical imaging of whole *Caenorhabditis elegans* and high-throughput detection of beta-amyloids in both mouse and human brain tissues associated with Alzheimer’s disease are demonstrated.

## 1. Introduction

Infrared (IR) absorption spectroscopy provides label-free, molecule-specific contrast by probing vibrational fingerprints. Extending this principle to imaging enables spatial mapping of chemical composition across materials and biological systems^1,2^. Recent developed quantum cascade laser (QCL)-based IR microscopes (**Fig. 1A**) have significantly improved the imaging throughput and widely used to localize spectral signatures^3-5^. However, because it relies on a focal plane array with a limited number of pixels, the field of view (FOV) decreases as the spatial resolution increases. In addition, the long wavelengths of mid-IR radiation restrict the diffraction-limited resolution to ∼3–30 µm^6^. Moreover, given to the strong water absorption, direct IR absorption measurements are difficult to perform in an aqueous environment, further hindering *in vivo* applications.

**Figure 1.**
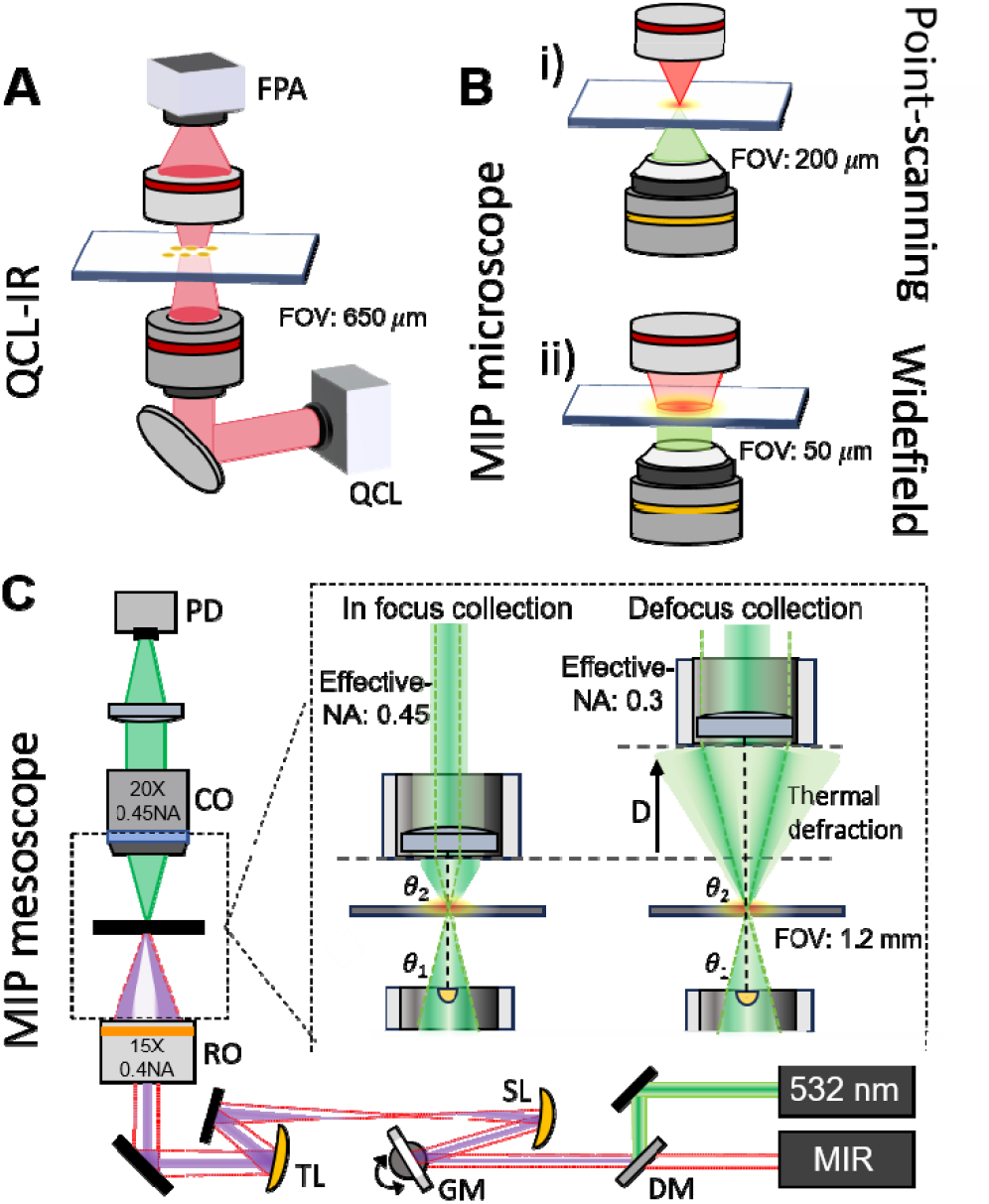
Comparison of QCL-IR microscope, MIP microscope and MIP mesoscope. (A) Schematic of QCL-IR microscope. IR illumination is provided by a QCL and the image of absorption is captured by a focal plane array (FPA). (B) Schematic of MIP microscope. IR pump beam and visible probe beam illuminate on the sample. In the point scanning mode, the photothermal image is formed by point scanning with sample stage. In the widefield mode, the photothermal contrast is acquired by subtracting the camera-captured frames between IR on and IR off status. (C) Schematic of MIP mesoscope. The IR pump beam and visible probe beam are combined and focused on the sample with a low-magnification objective lens. Imaging is performed by laser scanning via a pair of galvo mirror. Inset: illustration of defocused collection mechanism used in MIP mesoscope. QCL: quantum cascade laser. MIP: mid-infrared photothermal. PD: photodiode. CO: collection objective. RO: reflective objective. TL: tube lens. SL: scan lens. GM: galvo mirror. DM: dichroic mirror.

Recently developed mid-infrared photothermal (MIP) microscopy has bypassed the limitations encountered by conventional IR microscopy. By optically sensing mid-infrared absorption through a visible probe beam, MIP microscope has emerged as a powerful tool for chemical imaging with submicron spatial resolution^7,8^. Through indirect measurement of the thermal effect instead of direct measurement of IR photon attenuation, photothermal detection mitigates water background, allowing MIP imaging of living cells and small organisms cultured in a medium^9,10^.

Early implementations demonstrated high sensitivity by employing a photodiode as detector, but were constrained in throughput due to the slow stage scanning and acquisition. In addition, the stage travel range was restricted to 200 µm (**Fig. 1B**)^9,11^. The adoption of spatially multiplexed camera-based widefield detection further increased throughput by enhancing the imaging speed. In this schematic, photothermal contrast is acquired by subtracting the camera-captured frames between IR on and IR off status (**Fig. 1B**)^12-14^. However, a common CMOS sensor has a limited photon budget on the level of tens of thousands. Consequently, frame averaging is mandatory to resolve the small modulation depths^12^. In addition, the excitation fluence of weakly focused mid-IR illumination drops rapidly as the field of view (FOV) expands, diminishing photothermal signal and often requiring thousands of averages^12^. As a result, achieving high-throughput widefield MIP generally demands pump sources with high pulse energy^15^, which are less accessible than QCLs.

Recent point-scanning MIP imaging with commercially available QCLs has reached video rate by employing galvanometric beam steering^16^. In galvo scanning, fast galvanometric mirrors deflect a tightly focused laser spot across a sample. This scheme maintains the high sensitivity by tight focusing and increases the imaging speed by more than two orders of magnitude relative to stage scanning.

Importantly, the FOV in a galvo-scanning system can be scaled to millimeter scale by using imaging lens with long focal length while maintaining the speed and spatial resolution, analogous to two-photon (2P) mesoscopes used for whole-brain functional imaging^17,18^. Based on this concept, we report the first MIP mesoscope that achieves millimeter-scale FOV while preserving submicron resolution (**Fig. 1C**). Our design utilizes an all-reflective scanning system that eliminates chromatic aberrations from visible to mid-infrared. By using low-magnification, medium-NA objectives, we reach a FOV of 1.2 mm with 650 nm spatial resolution. An optimized defocused collection scheme is designed for amplifying photothermal signal and further maximizing the usable FOV. With the developed MIP mesoscope, we demonstrated *in vivo* chemical imaging of whole *Caenorhabditis elegans* and achieved more than 3-orders-of-magnitude higher-throughput compositional mapping of β-amyloid in mouse and human brain tissues with Alzheimer’s disease^19^, substantially advancing tissue-level studies, which is essential for systematic and statistically robust investigations of molecular pathology on organ level.

## 2. Results

### 2.1 MIP mesoscope and performance characterization

As shown in the **Fig.1C**, the MIP mesoscope adopts a co-propagation scheme, where the mid-infrared pump and the visible probe are combined and focused onto the sample by the same objective lens. Mid-IR excitation is provided by a QCL outputting pulses at 600 kHz with 300 ns pulse duration, which induces a transient thermal modulation. The IR beam is combined with a 532-nm continuous-wave visible probe beam via a dichroic mirror. The combined beam is scanned by an X–Y pair of galvanometric mirrors. A pair of concave mirrors relay the scanned beam to the objective back pupil of a 15×, 0.45-NA reflective objective, providing 2.5× beam diameter to fill the aperture. The all-reflective scanning relay eliminates chromatic aberrations, enabling IR spectroscopic imaging without re-alignment across visible and IR wavelengths. The probe, carrying the photothermal modulation, is collected by a 20X, 0.45 NA refractive objective. The imaging performance of designed system is simulated by Zemax as shown in **Fig. S1**. Notably, the spatial resolution of our system is determined by the diffraction limit of the visible beam, which is 667 nm through an objective 0.4 NA.

In our MIP mesoscope, the photothermal signal is generated by a defocusing collection scheme as shown in inset of **Fig.1C**. In this scheme, the NA mismatch introduced between the focusing and collection lenses converts the thermal deflection into an intensity change at the detector. Compared with the pinhole/iris spatial filtering used in classic MIP implementations^9^, the defocused propagation preserves the full usable field of view of the reflective objective (**Fig. S2**). The defocused collection scheme in our system serves two important functions. First, it introduces a numerical aperture (NA) mismatch between the focusing and collection objectives (**Fig. S3**), which enhances the photothermal-induced intensity modulation at the detector, thereby improving the photothermal signal contrast. Second, the defocus enlarges the effective FOV of the collection system, enabling us to fully exploit the wide-FOV advantage of the reflective objective.

With the MIP mesoscope constructed (**Fig. S4)**, we tested its performance from four perspectives: spectral fidelity, spatial resolution, FOV, and sensitivity. To validate spectral fidelity, we performed hyperspectral imaging of 1-μm diameter polystyrene (PS) beads. MIP spectra were acquired by sweeping the QCL from 1400 to 1650 cm^−1^ in 5 cm^−1^ steps and collecting an MIP image at each wavelength. The spectrum from single PS particle was normalized by the power spectrum of QCL measured and shown in **Fig. 2A**. Compared with conventional FTIR reference (**Fig. S5**), the spectrum is in good agreement in peak positions and relative intensities throughout the fingerprint region. The spatial resolution of the system is evaluated at the image taken at 1450 cm^−1^ (**Fig. 2B**) by selecting a single bead. The point spread function is measured with full width at half maximum (FWHM) of 0.65 μm in the x direction and of 0.86 μm in the y direction after deconvolution with the diameter of the bead. To characterize the effective FOV, we imaged 3-μm diameter polymethyl methacrylate (PMMA) beads at their absorption peak at 1729 cm^-1^ (**Fig. 2C**). The mesoscope provides a FOV of 1.2 mm × 1.2 mm. Illumination uniformity was quantified from horizontal and vertical line profiles drawn across the image (**Fig. 2D**). The normalized intensity varies by only 1% from center to edge along both axes, indicating highly uniform excitation and collection across the full field. Lastly, we characterized the system sensitivity by performing hyperspectral imaging of 500-nm diameter PMMA beads. The transmission image and MIP images at various wavenumber are shown in **Fig. 2E**. The hyperspectral stack is acquired by sweeping the QCL from 1400 to 1780 cm^-1^ at 5 cm^-1^ intervals. The spectrum of an individual bead is shown in **Fig. 2F**, agrees with the FTIR reference shown in **Fig. S5**, further validating the high sensitivity of the built MIP mesoscope.

**Figure 2.**
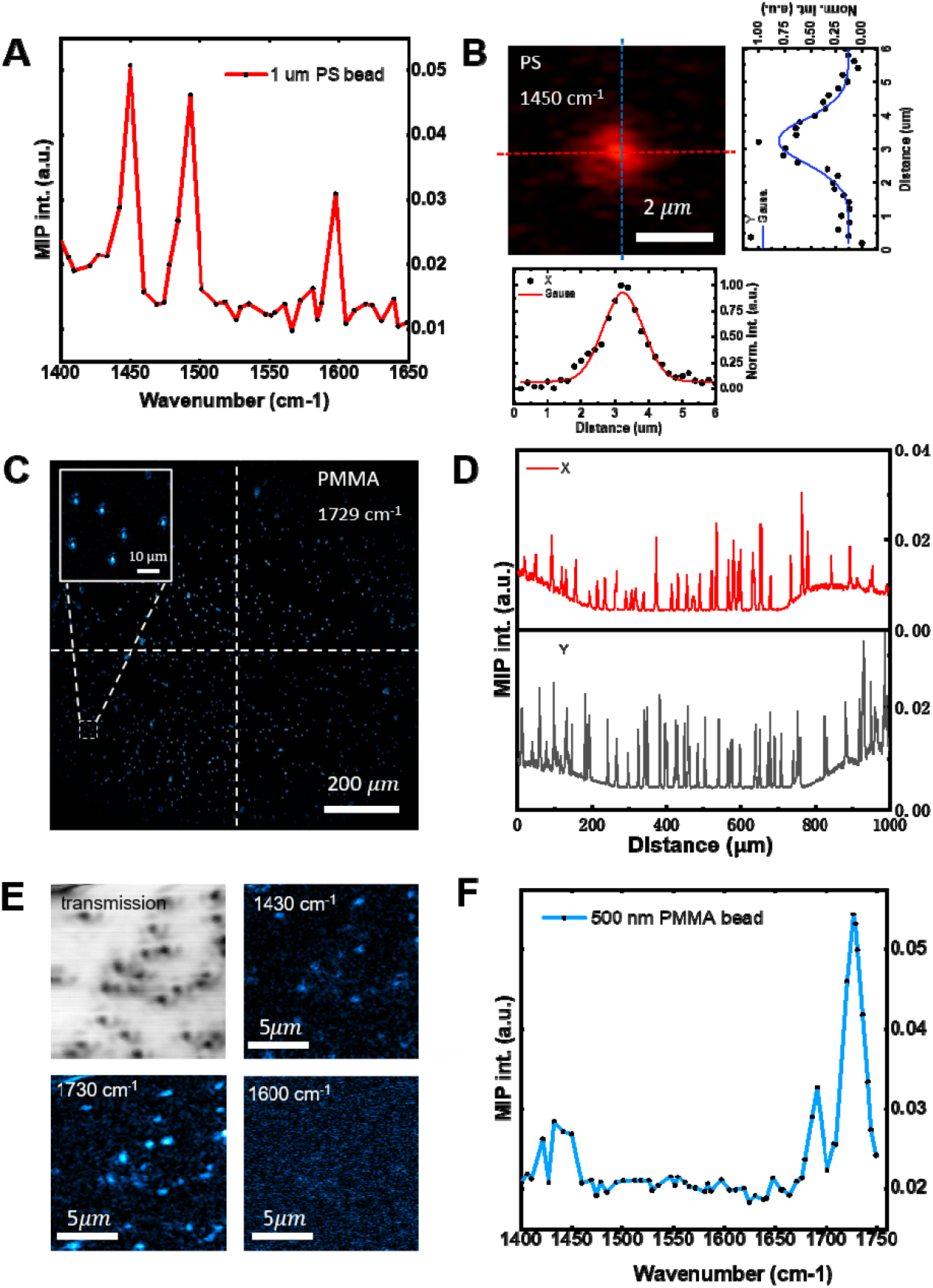
MIP mesoscope performance characterization. (A) Spectral fidelity evaluated using 1 µm polystyrene (PS) beads. (B) Resolution test by intensity profiles across the center of a selected bead imaged at 1450 cm^-1^ in horizontal and vertical directions, with Gaussian fitting used to determine the full width at half maximum (FWHM), yielding FWHM_vertical_ = 1.19 µm and FWHM_horizontal_ = 1.32 µm. (C) Field-of-view (FOV) test using 3 µm PMMA beads imaged at 1729 cm^-1^. (D) Uniformity assessed by plotting horizontal and vertical intensity profiles across the FOV. (E) Multi-wavenumber MIP imaging of 500 nm PMMA beads. Transmission image is also shown as a reference. (F) Sensitivity demonstration by spectral fidelity of 500 nm PMMA beads.

### 2.2 Chemical imaging of whole *C. elegans in vivo*

*Caenorhabditis elegans* (*C. elegans*) was the first multicellular organism to have its genome fully sequenced. It contains ∼20,000 genes, ∼60–80% of which are homologous to human gene^20^. Chemical imaging of *C. elegans* by stimulated Raman scattering (SRS) microscopy allowed researchers to examine the role of fat in aging^21^. However, the FOV of SRS microscopy is insufficient to cover the entire worm. IR spectroscopy could provide a larger field of view and is more sensitive in the fingerprint window. Yet, IR spectroscopic imaging of *C. elegans* in native, aqueous environments remain challenging. Traditional IR imaging is only applicable to dried sample, and its limited spatial resolution cannot resolve chemistry at the level of individual organs. MIP microscopy overcomes the above-mentioned issue and enabled the live worm imaging^16,22^, but the restricted FOV only captures part of the worm, requiring mosaicking multiple images to capture a whole worm.

Our MIP mesoscope, with an extended FOV, is able to capture entire *C. elegans* within tens of seconds, enabling *in vivo* acquisition of living dynamics with chemical information from the intact organism (see **Movie S1**). We imaged worms at 1550 cm□^1^ by targeting the amide II band of proteins as shown in **Fig. 3A**, and 1740 cm□^1^ by targeting esterified lipid as shown in **Fig. 3B**. The merged image (**Fig. 3C**) reveals the spatial distribution of protein and esterified lipid across the worm body. With submicron spatial resolution, the fine features of various organs are visible (**Fig. 3D)**. Including the (i) densely packed lipid droplets (LDs) in epidermis. (ii) an embryo rich in lipid, (iii) the head region with prominent LDs and a modest protein signal, and (iv) the tail region, which is protein-rich and exhibits an annular lipid feature.

**Figure 3.**
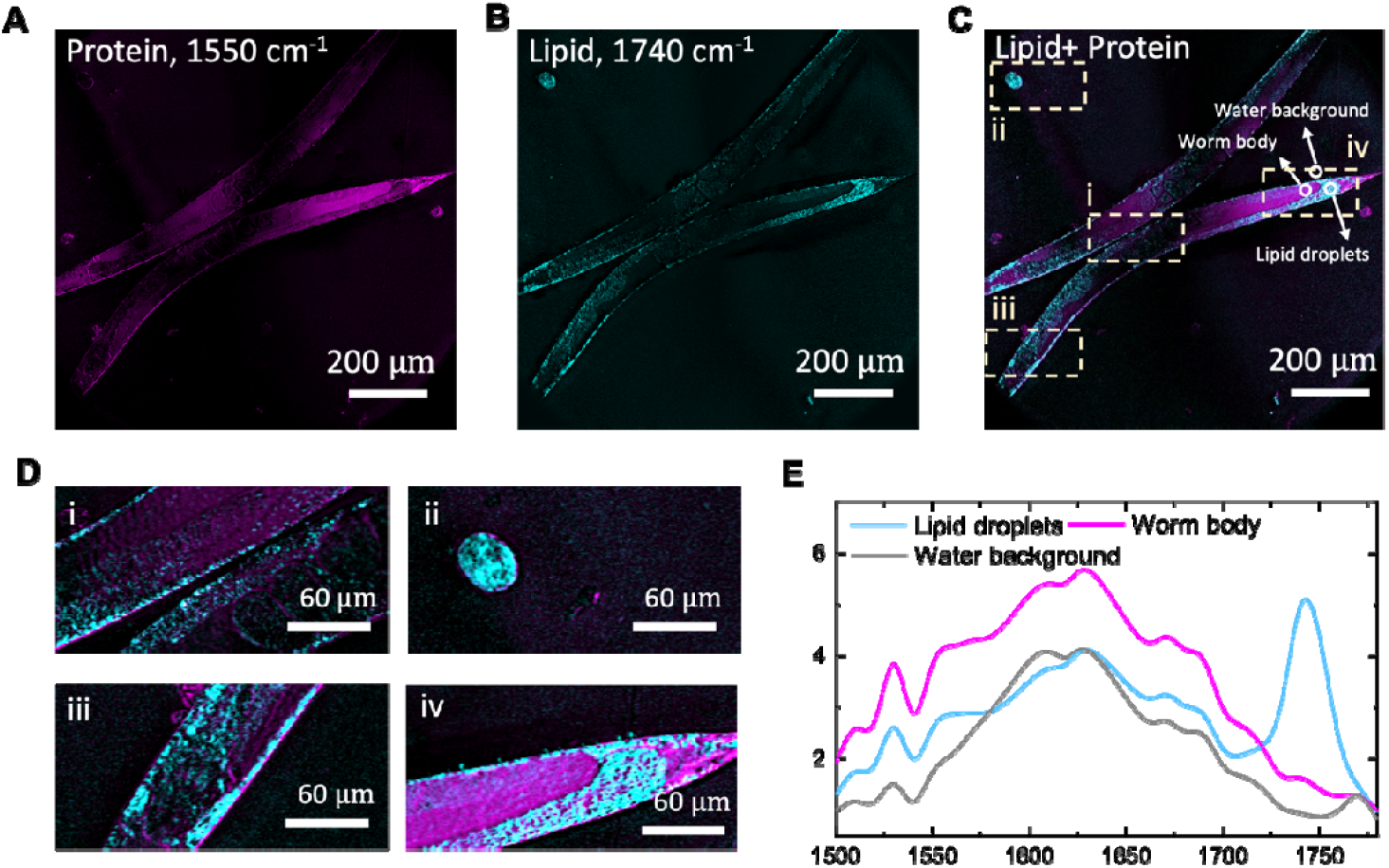
Spectroscopic imaging of living *C. elegans*. (A) Imaging protein of worms acquired at 1550 cm□^1^. (B) Imaging lipid of the same worm acquired at 1740 cm□^1^, highlighting lipid droplets (LDs). (C) Merged image of (A) and (B), showing the spatial distributions of proteins and lipids within the worm body. (D) Magnified views of selected regions: (i) densely packed lipid droplets (LDs) in epidermis and the developing embryos, (ii) an embryo rich in lipids, (iii) the head region with prominent LDs and a modest protein signal, and (iv) the tail region, which is protein-rich and exhibits an annular lipid feature. (E) Representative spectra extracted from three regions of interest (ROIs) marked in (C): LD-rich, protein-rich, and aqueous background. The image shown in panel A and B took 20 seconds with a pixel dwell time of 20 μs.

To obtain localized molecular spectra, we performed hyperspectral imaging of the whole worm by sweeping the QCL from 1500 to 1780 cm□^1^ in 10 cm^-1^ increments. The spectra were normalized to the measured QCL power at each wavenumber. The spectra of three regions of interest (ROIs) with LD-rich, protein-rich, and aqueous background (circles in **Fig. 3C**) respectively were show in **Fig. 3E**. The LD-rich region shows a pronounced peak near 1740 cm□^1^, whereas the protein-rich region exhibits stronger signal near 1550 cm□^1^. The background exhibits a broad band centered at around 1650 cm□^1^ attributable to the H–O–H bending mode of water. With the ability for imaging molecules in whole worm at sub-micron resolution *in vivo*, the developed MIP mesoscope provides a practical tool for investigating the developmental dynamics and disease-related processes in *C. elegans*.

### 2.3 High-throughput chemical histology of protein secondary structure

In addition to *in vivo* study, the MIP mesoscope can be used to boost the throughput in chemical histology by more than three order magnitude^19^. Here, using tissues affected by Alzheimer’s disease, we reveal the protein secondary structure changes in brain tissues from both mouse models and human patient samples.

We first imaged brain sections from a 6-month-old 5xFAD transgenic mouse which developed pronounced β-amyloid (Aβ) deposits in the hippocampus and cortex^19^. We confirmed the Aβ enrichment by colocalized fluorescence image as shown in **Fig. S6**. Two MIP images at 1630 cm□^1^, where β-sheet dominant, and 1658 cm□^1^, where α-helix is dominant were taken subsequently. The whole 5xFAD mouse brain map is acquired by stitching 6 by 13 images, as shown in **Fig. 4A**. To visualize the β-sheet enrichment, a ratio image I□□□□/I□□□□ was then computed as shown in **Fig. 4B. Fig. 4C** is the zoom-in view of the boxed regions in Fig. 4B, where the hotspots are visualized, indicating Aβ plaques. Age-matched wild-type mice served as the control group. The whole transmission image was acquired by stitching 12 by 12 images as shown in **Fig. 4D** and the corresponding I□□□□/I□□□□ ratio image was then generated (**Fig. 4E**). In contrast to the 5xFAD group, zoom-in views of the boxed regions (**Fig. 4F**) confirmed the absence of β-amyloid plaques.

**Figure 4.**
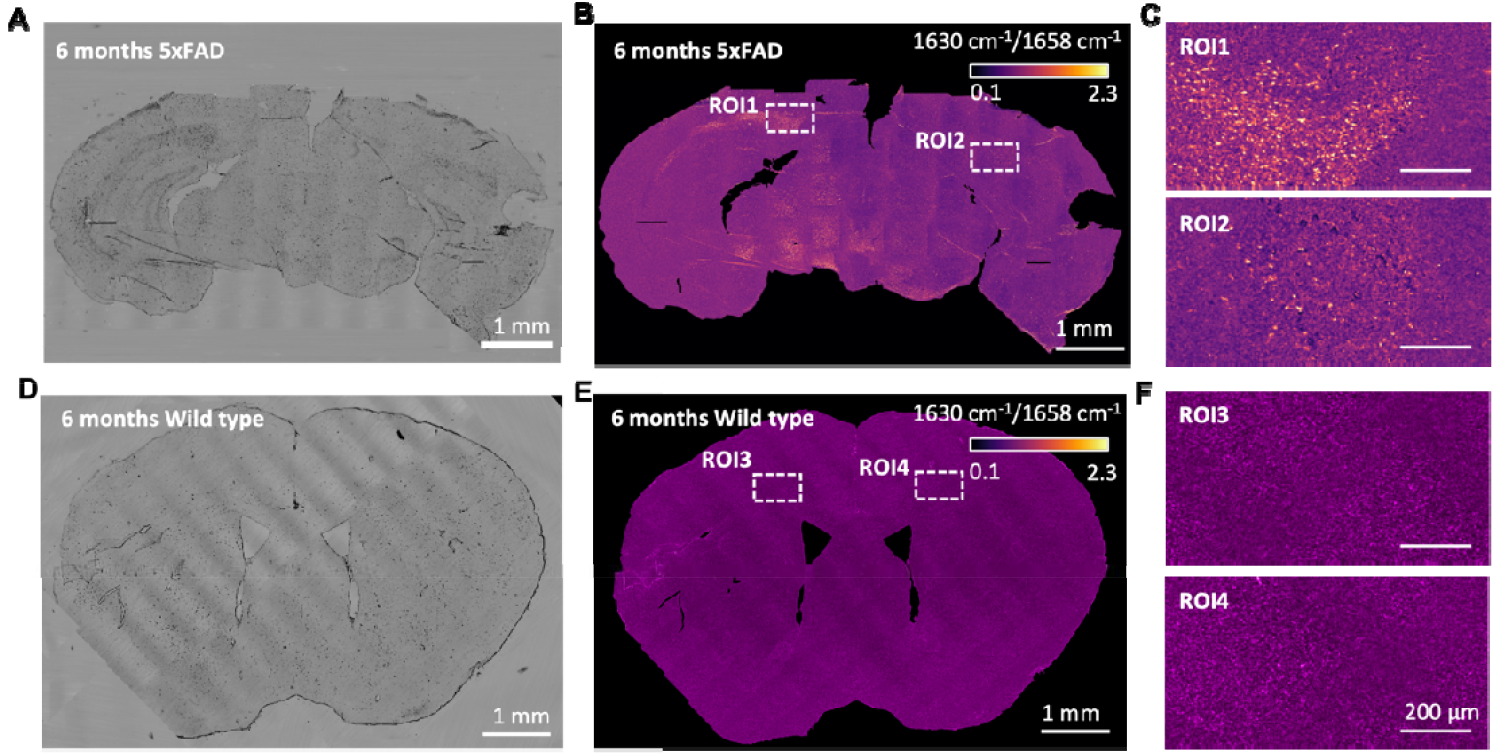
Mapping protein secondary structure in whole mouse brain. (A) Transmission image of 5xFAD mouse brain tissue. (B) Ratio map of the tissue calculated from the intensity ratio at 1632/1658 cm□^1^, with the boxed area indicating β-amyloid–rich regions. (C) Zoom-in views of the boxed regions in (B), where bright spots represent β-amyloid plaques. (D) Transmission image of wild-type mouse brain tissue. (E) Corresponding ratio map calculated at 1632/1658 cm□^1^, showing no obvious β-amyloid–rich regions. (F) Zoom-in views of the boxed regions in (E), confirming the absence of β-amyloid–rich structures. The whole brain image took ∼40 minutes at each wavenumber, with a pixel dwell time of 20 μs.

We next acquired hyperspectral images of the 5xFAD tissue with a finer spectral resolution, spanning 1500–1720 cm□^1^ in 2 cm□^1^ increments. The transmission image of the selected region is shown in **Fig.□5A**. Using two-color composite images at 1630□cm□^1^ (magenta) and 1658□cm□^1^ (cyan) (**Fig□5B**), β□sheet–enriched regions appear as bright spots. Zoom in views of representative β□sheet–enriched aggregates from **Fig.□5B** are presented in **Fig.□5C**. Spectra were extracted from five normal protein ROIs (circled in **Fig.□5B**) and five aggregates (highlighted in **Fig.□5C**), and are shown in **Fig.□5D** and **5E**, respectively. Compared with non□enriched ROIs, β□sheet–enriched ROIs exhibit a pronounced peak near 1632□cm□^1^. The averaged spectra (mean of five ROIs per group), shown in **Fig.□5F**, highlight the secondary structure changes in protein within aggregate□rich regions.

**Figure 5.**
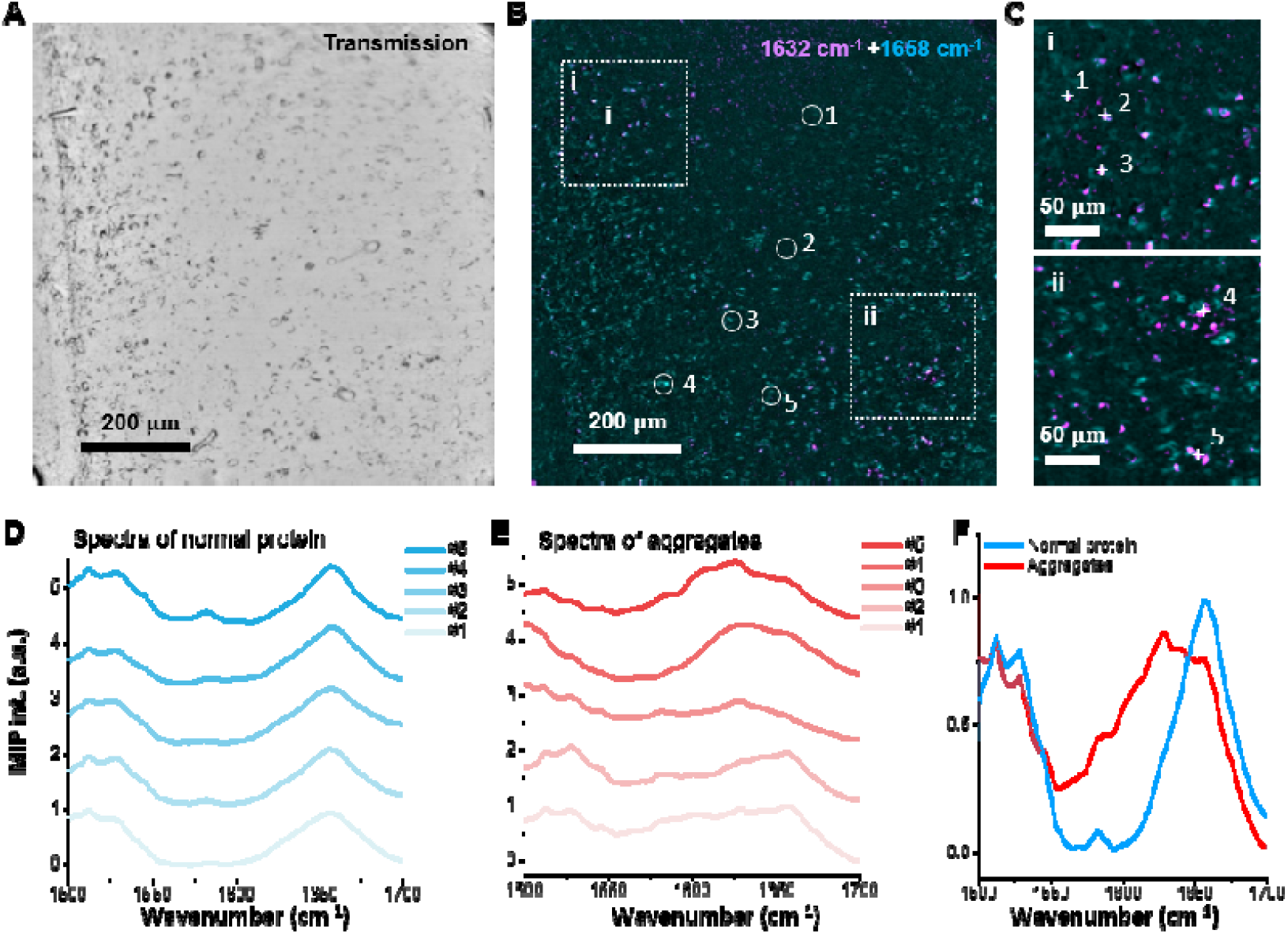
MIP spectroscopic imaging of mouse brain tissue. (A) Transmission image of the sample. (B) Merged MIP image at 1632 cm□^1^ and 1658 cm□^1^ of the same sample, revealing distinct spatial distributions of β-amyloid and normal proteins. (C) Zoomed-in view of the region of interest. (D) Representative spectra from five points in panel (C) corresponding to β-amyloid aggregates. (E) Representative spectra from five regions of interest (ROIs) corresponding to normal protein areas. (F) Average spectra of the selected aggregates and normal protein ROIs, showing clear spectral differences between the two.

We further examined brain tissues from postmortem AD patients. Fig. 6A shows a photograph of the paraffin embedded human brain tissue sample mounted on a histology glass slide together with a bright-field microscopic image that illustrates the typical sample size. **Fig. 6B** presents an immunofluorescence image of the same tissue after performing the spectroscopic imaging, clearly revealing the presence of amyloids in tissue and vessels. The bright field image of a vessel is shown in **Fig. 6C**. A two-color composite of MIP images at 1632 cm□^1^ and1658 cm^-1^ is shown in **Fig. 6D**. We performed spectroscopic imaging of the sample from 1500-1720 cm^-1^ in 2 cm^-1^ steps. Then regions of interest, including five areas with elevated β-sheet content and three regions with predominantly native protein fold, were selected for spectral comparison (Fig. 6E). Compared with the native protein regions (ROI 1–3), the selected areas (dots 1–5) showed markedly higher intensity at 1632 cm□^1^, indicating the presence of β-sheets corresponding to aggregated Aβ. Importantly, the larger field of view provides two major advantages: first, it enables direct comparison of multiple amyloid deposits within the same section; second, larger area allows more precise registration with other downstream imaging modalities, such as spatial omics. Such comprehensive investigation was not feasible with previous techniques, which were restricted to small regions of interest (typically 100 100 μm).

**Figure 6.**
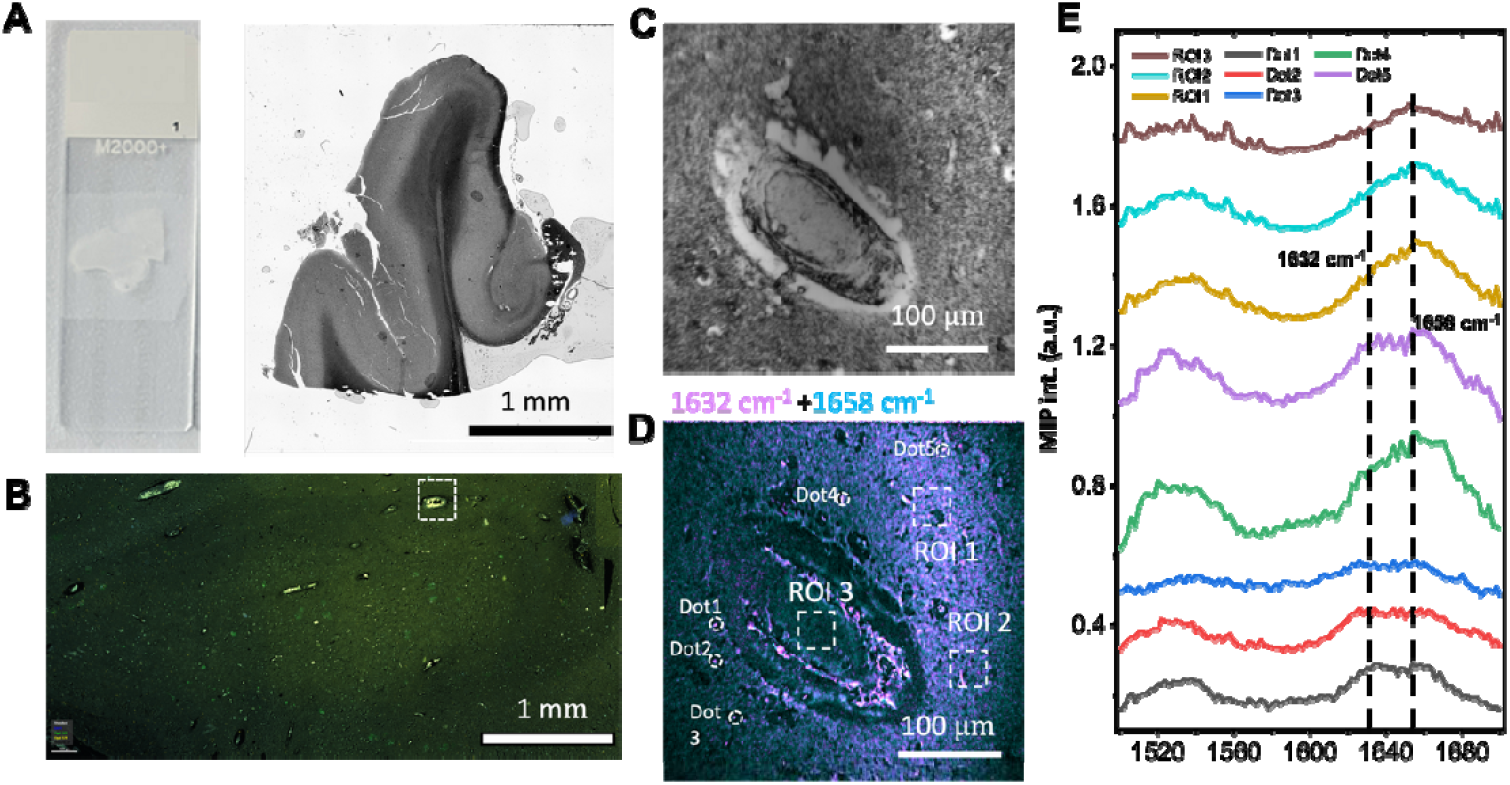
MIP spectroscopic imaging of human brain tissues. (A) Photograph of a histological glass slide with human brain tissue together with a bright-field image showing the typical size of the sample. (B) Immunofluorescence image of the same tissue, clearly revealing a representative vessel containing amyloid deposits. (C) Transmission image of a vessel in human brain tissue section. (D) Merged MIP image at 1632 cm□^1^ and 1658 cm□^1^ of the same sample with selected regions of interest (ROIs) highlighted. (E) Spectra obtained from five points and three ROIs indicated in panel (D). The MIP image took 1.6 seconds at each wavenumber, with a pixel dwell time of 10 μs.

## 3. Discussion

By leveraging an all-reflective laser scanning architecture, low-magnification with medium-numerical-aperture objectives, and a defocused collection scheme, we demonstrated a MIP mesoscope and achieved a 1.2 × 1.2 cm^2^ field of view while maintaining submicron spatial resolution. This capability enables *in vivo* chemical imaging of entire *C. elegans* in a single scan, allowing direct observation of dynamic processes throughout the organism. In addition, high-throughput detection of β-amyloids in both mouse and human brain tissues associated with Alzheimer’s disease were demonstrated.

The illumination uniformity in our current setup can be further improved. Our scanning system employs a 4-f optical relay using concave mirrors only. Illumination non-uniformity is observed in images with maximum FOV **(Fig. S7)**. Using Zemax simulations, we explored a modified configuration in which a convex mirror is added after the 4f system to correct aberrations **(Fig. S8)**. The simulated irradiance maps show that this modification significantly reduces the intensity variation between the image center and periphery.

Beyond relay optimization, further enhancement of illumination uniformity as well as resolution could be achieved through freeform optical design of the focusing objective^23^. The reflective objective currently employed is a commercial one with Cassegrain design. Such a design unavoidably results in non-uniform illumination across the FOV due to central block, which in turn limits image quality. In addition, the medium NA of this objective constrains the achievable resolution of our system. These limitations could be overcome by employing a custom-engineered freeform all-reflective objective, which would allow optimized aberration correction and more uniform light delivery across the large FOV.

The reported MIP mesoscope addressed the limited FOV issue in MIP microscopy. Conventional MIP microscope has been successfully applied in diverse biological studies ranging from fingerprinting single viruses^24,25^, bacteria^26,27^ to cells^28^. However, because of the relatively small FOV, the applications in large specimens such as brain tissues and whole animals such as *C. elegans*^9^ are largely restricted. MIP mesoscopy opens broad opportunities in biomedical research, such as label-free mapping of microplastics in tissues, high-throughput cell imaging, quantitation of protein aggregations in neuro-degenerative disorders, and mapping of intracellular metabolites in cancer tissues and integration with spatial omics for correlative molecular analysis^29^.

Such fast-imaging capability enables scanning of multiple adjacent sections and reconstruction of amyloid structures within the 3D architecture of complex organs like the brain. Volumetric mapping will be crucial for revealing how pathological features — such as tissue responses to amyloid deposits — are spatially organized within the tissue. This approach opens new possibilities to study structural heterogeneity and tissue responses in unprecedented detail, which has so far been limited by slow imaging speed. We anticipate that this optical mesoscopy modality will integrate with and enrich the broader omics landscape, opening new opportunities for tissue-level biomedical discovery.

## 4. Materials and Methods

### PMMA and PS particles

The PMMA and PS particles in solution form were first diluted with deionized water. Around 2 μl of the solution was then dropped on the surface of a calcium fluoride (CaF2) substrate with a thickness of 0.2 mm for imaging.

### *C. elegans* maintenance and mounting

N2 worms were cultured on nematode growth medium (for 1L: phosphate buffer 20mM, CaCl2 0.8mM, MgSO4 0.8mM, agar 17.5g, NaCl 3.0g, peptone 2.5g, cholesterol 0.005g) and grown at 15°C. Plates were seeded with OP-50 Escherichia coli. For imaging, young adult worms (1-3 days old) were mounted via picking into 1% NaN3 in M9 buffer on a 24x60mm #1 thickness cover slide. A calcium fluoride (CaF2) disk was then lowered onto the worms and fixed in place with tape.

### Mouse brain tissue

The 5xFAD mouse is a widely used transgenic model for studying amyloidosis in Alzheimer’s disease (AD). These mice carry human APP and PSEN1 genes with five AD-related mutations: three in APP (Swedish K670N/M671L, Florida I716V, and London V717I) and two in PSEN1 (M146L and L286V). A characteristic feature is the selective accumulation of intraneuronal amyloid-β (Aβ) in the cortex^30^.

By 12 months of age, the mice develop abundant extracellular amyloid plaques with well-defined fibrillar cores, providing a robust system for optical and spectroscopic imaging studies. For the experiment, mice were transcardially perfused with 4% paraformaldehyde (PFA) in PBS, and brains were carefully dissected. Post-fixation was performed in 4% buffered formalin at 4 °C. The tissue was then sectioned at 16 µm thickness, mounted on standard histological glass slides, and air-dried prior to measurements.

All animal experiments were conducted in accordance with the requirements of the Ethical Committee of Lund University (Malmö/Lund djurförsöksetiska nämnd 11908/2019-16). 5xFAD mice (hAPPswe, PSEN1dE985Dbo/Mmjax) were obtained from Jackson Laboratories (USA), and the presence of the human APP695 transgene was verified by PCR.

### Human brain tissue

N1792 tissue, previously characterized and reported^31^, was used in this study. All procedures complied with relevant ethical guidelines and were approved by BRAIN UK (UK Brain Archive Information Network) under REC reference 19/SC/0217.

Formalin-fixed paraffin-embedded (FFPE) sections were deparaffinized and imaged using two complementary approaches: the MIP mesoscope for large-area infrared mapping and the Mirage optical photothermal infrared (O-PTIR) system (Photothermal Corp., Santa Barbara, USA) for spectroscopic validation. Following spectroscopic measurements, the sections were rehydrated through an ethanol series and stained to confirm amyloid presence. According to the manufacturer’s protocol, Amytracker® 520 (1:1000; Ebba Biotech, Solna, Sweden) was applied together with DAPI (1:10,000; Sigma) for 30 min at room temperature, then mounted with DAKO medium. Imaging was performed using a BioTek Cytation 5 Cell Imaging Multi-Mode Reader and a Zeiss Axio epifluorescence microscope system.

### Image processing

All MIP images were imported to ImageJ (National Institutes of Health) for analysis. For the beads imaging for resolution test, the intensity profiles were plotted at the center of the bead in the horizontal and vertical directions. Gaussian fitting was used to determine the FWHM of the beads. For illumination fidelity, the intensity profiles were plotted at the center of the FOV in the horizontal direction.

For the *C. elegans* images shown in Fig. 3A, we used the ImageJ image calculator to remove the water background. In detail, we used the image at 1600 cm□^1^, which contains a weak protein contribution but relatively strong water absorption, was used together with the image at 1550 cm□^1^, which represents the protein signal but also contains water background. A background-corrected image highlighting the protein distribution was obtained by calculating *I*□□□□ *– I*□□□□ *× (*⟨*I*□□□□⟩*/*⟨*I*□□□□⟩) on a pixel-by-pixel basis, where ⟨I⟩ denotes the mean intensity of the corresponding background. Spectra were taken with a step of 5 cm^-1^ and corrected by the IR power intensity measured.

For large-area MIP imaging of mice brain tissues, we employed a multi-scale scanning system that synchronously controlled a laser-scanning module and a precision motorized translation stage (MCM3000, Thorlabs). The system first performed high-resolution imaging of a local region measuring 800 × 800 μm^2^ using laser scanning. Subsequently, the translation stage moved the sample to an adjacent region with a step size of 0.6 mm and a 20% overlap for the next scan. A serpentine scanning strategy was implemented to optimize the scan path: subregions in odd-numbered rows were scanned from left to right, while those in even-numbered rows were scanned in the reverse direction from right to left. This approach minimized idle travel distances and improved overall scanning efficiency.

For spectra of brain tissues, we performed hyperspectral imaging of the brain tissue by sweeping the QCL from 1500 to 1780 cm□^1^ in 2 cm^−1^ increments. The spectra were normalized to the measured QCL power at each wavenumber.

## Data availability

All experimental data and any related experimental background information not mentioned in the text are available from the authors upon reasonable request.

## Acknowledgements

This work is partially supported by R44GM154516 to JXC.

## Author Contributions

J.Y. and J.-X.C. initialized the idea of MIP mesoscope; R.T. and J.Y. designed and built the MIP mesoscope setup and carried out the experiments; R.T. and J.Y analyzed the experimental results. R.T. performed the simulation. B.W. performed all worm culture and mounting, as well as editing the manuscript. O.K. and D.G. provided all the brain tissue samples and edited the manuscript. H.L. helped data processing of whole mouse brain tissue. J.L. provided some human brain images. J.-X.C. supervised the research and the development of the manuscript. R.T. wrote the manuscript with input from all authors.

