## Supplemental figures for "Mid-Infrared Photothermal Mesoscopy with Millimeter Field of View and Sub-micron Spatial Resolution"

**Supplementary Information**

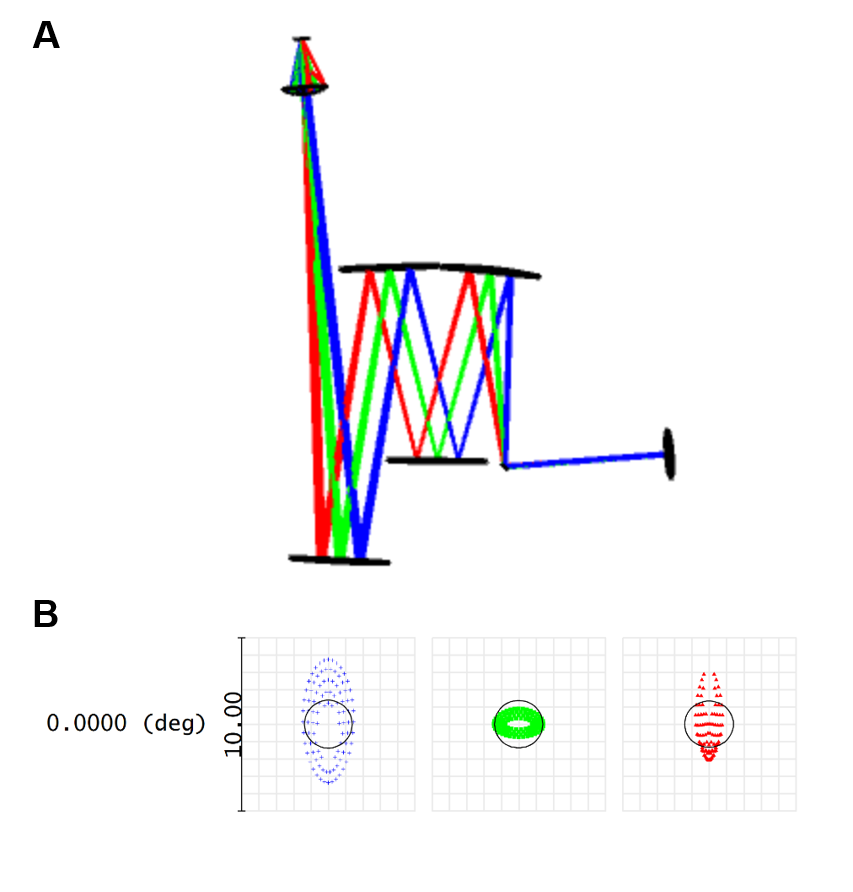

**Figure S1. Zemax simulation of the focusing optical path in the MIP mesoscope.** (A) Simulated optical path of the scanning system in the current MIP mesoscope. The reflective optical design enables large field-of-view laser scanning while minimizing chromatic aberrations. (B) Simulated imaging quality of the scanning system of current MIP mesoscope. Spot diagrams at different input angle shows the residual astigmatism and aberrations still existing.

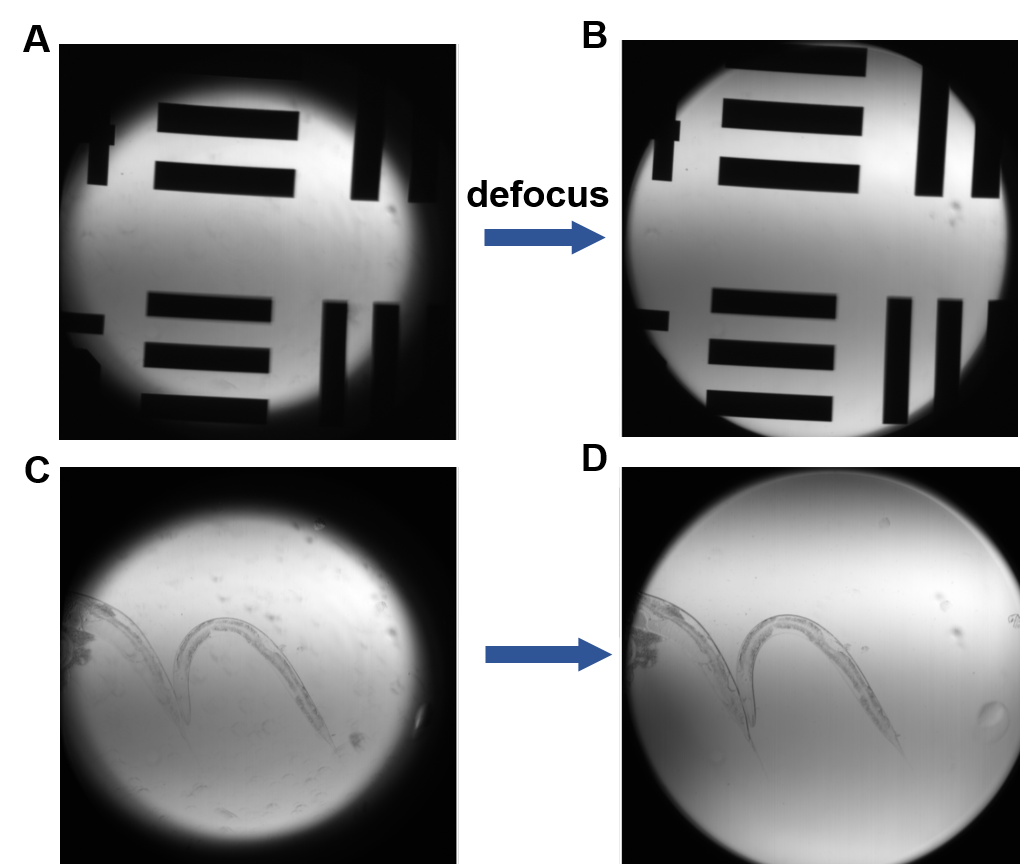

**Figure S2. Effect of defocusing the collection objective on field of view (FOV).** (A) Resolution target image acquired with the collection objective in focus. (B) Same region as (A) with the collection objective defocused, showing an enlarged FOV. (C) *C. elegans* image acquired with the collection objective in focus. (D) Same region as (C) with the collection objective defocused, also exhibiting an enlarged FOV.

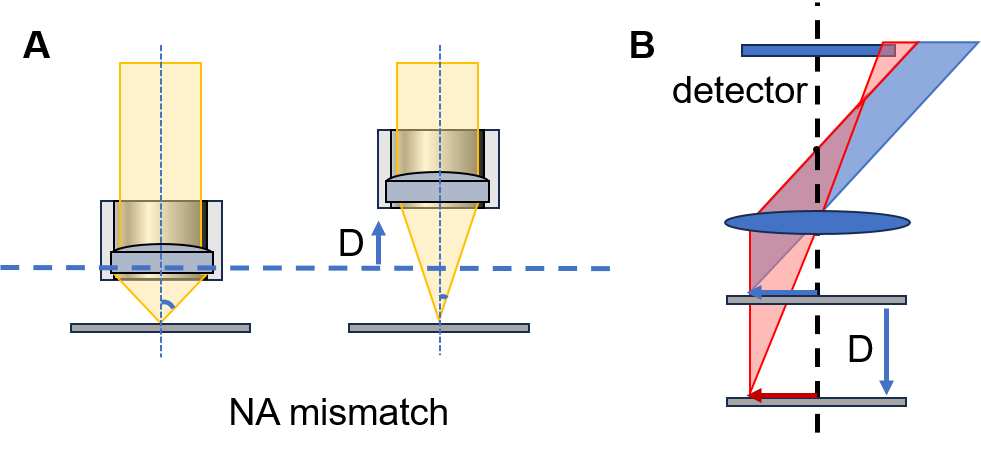

**Figure S3. Illustration of NA mismatch and its role in enlarging the field of view (FOV).** (A) Numerical aperture (NA) is defined as NA = n·sinθ. Increasing the working distance reduces the NA of the collection objective by decreasing the collection angle θ. (B) Schematic showing in-focus (blue) and defocused (red) light paths. For the same FOV, defocused rays can still be collected by the detector, effectively enlarging the collection FOV.

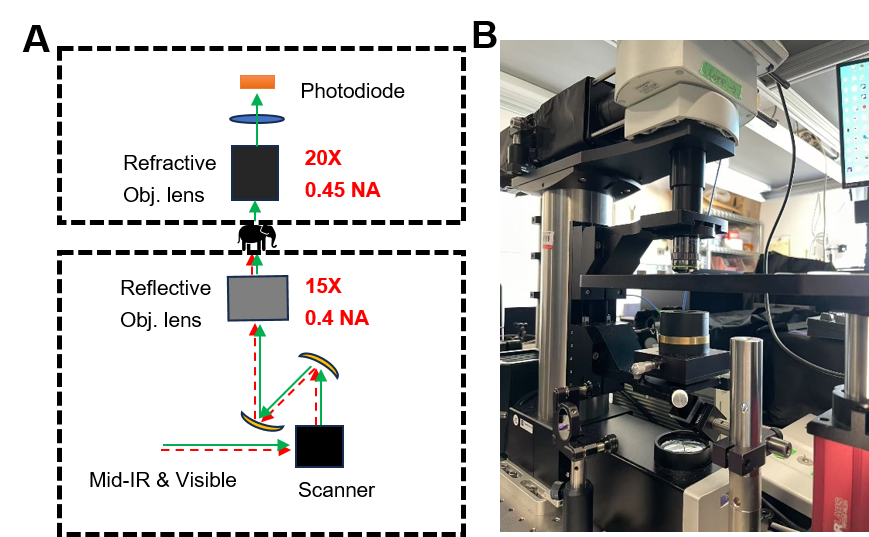

**Figure S4. The constructed MIP mesoscope.** (A) Simplified schematic of the MIP mesoscope, showing the 15× reflective objective (0.4 NA) for excitation and the 20× refractive objective (0.45 NA) for signal collection. (B) Photograph of the constructed MIP mesoscope system.

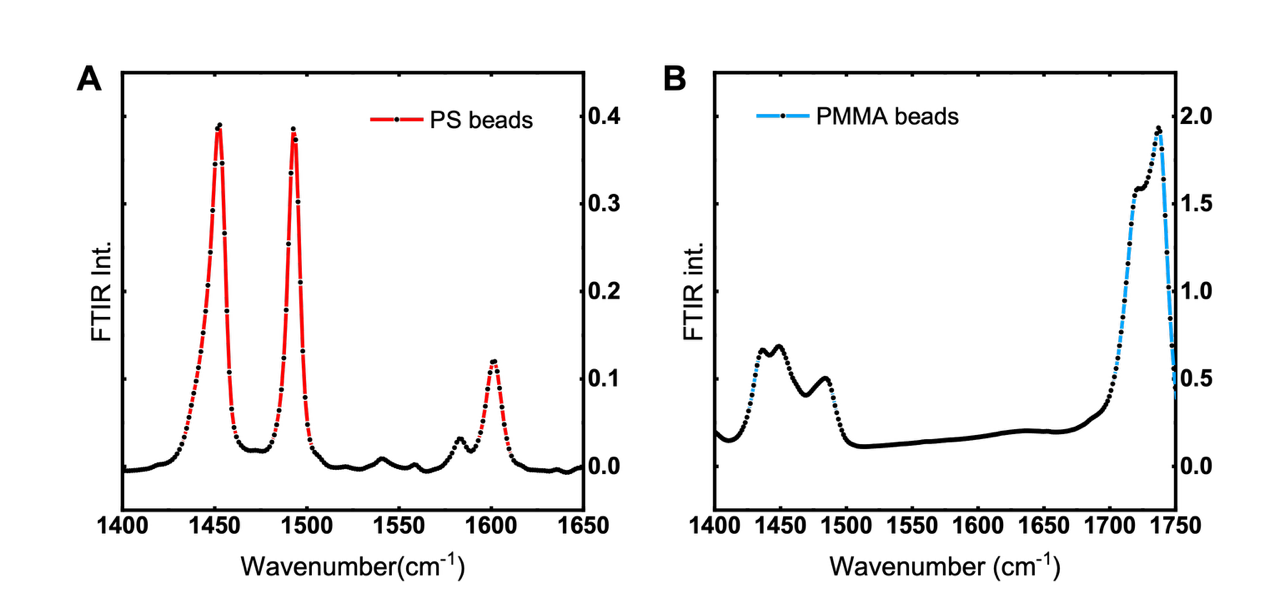

**Figure S5. FTIR spectra.** (A) FTIR spectrum of PS. (B) FTIR spectrum of PMMA. The spectra were recorded by Nicolet FT-IR with ATR.

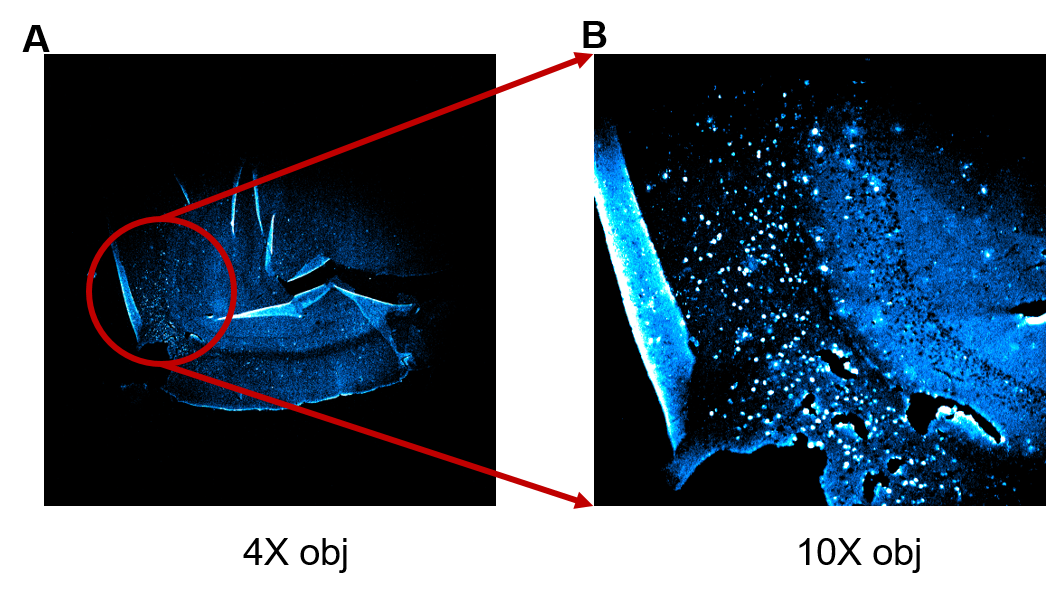

**Figure S6. Fluorescence microscopy confirming the enrichment of Aβ in 5xFAD brain tissue.** (A) Fluorescence image of a 5xFAD sample acquired with a 4× objective, showing aggregated Aβ deposits. (B) Magnified view of the circled region in (A) acquired with a 10× objective, where individual Aβ aggregates are more clearly resolved.

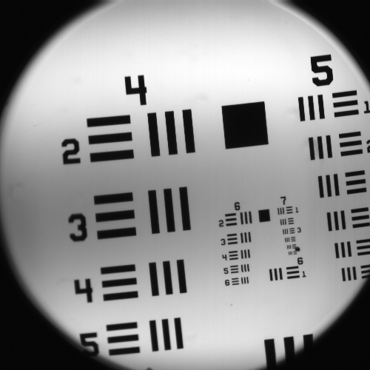

**Figure S7. Illumination profile of the MIP mesoscope.** Uneven illumination was observed in the current setup, with a darker intensity region at the center.

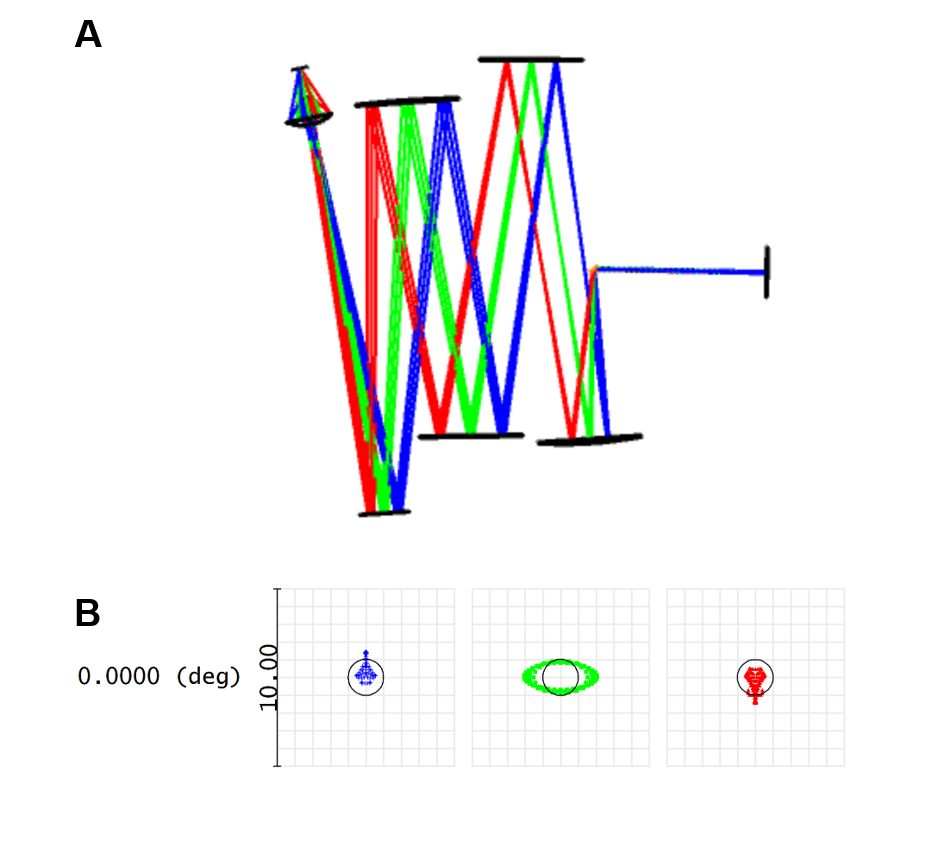

**Figure S8. Zemax simulation of the focusing optical path in the modified configuration.** (A) Zemax simulation of the modified configuration, where a convex mirror was added after the 4f system to reduce aberrations. (B) Spot diagrams showing improved imaging quality, particularly at the periphery of the field.
